# Can higher aggressiveness effectively compensate for a virulence deficiency in plant pathogen? A case study of *Puccinia triticina*’s fitness evolution in a diversified varietal landscape

**DOI:** 10.1101/2023.06.09.544363

**Authors:** Cécilia Fontyn, Kevin JG Meyer, Anne-Lise Boixel, Corentin Picard, Adrien Destanque, Thierry C Marcel, Frédéric Suffert, Henriette Goyeau

## Abstract

Plant resistances impose strong selective pressure on plant pathogen populations through the deployment of resistance genes, which leads to the emergence of new virulences. The pathogen adaptation also involves other parasitic fitness traits, especially aggressiveness components. A previous study on Puccinia triticina, the causal agent of wheat leaf rust, revealed that the distribution frequency of virulences in the French pathogen population cannot be fully explained by the major resistance genes deployed in the landscape. From 2012 to 2015, two dominant pathotypes (distinguished by their combination of virulences) were equally frequent despite the theoretical advantage conferred to one pathotype (166 317 0) by its virulence to Lr3, frequent in the cultivated landscape, whereas the other (106 314 0) is avirulent to this gene. To explain this apparent contradiction, we assessed three components of aggressiveness — infection efficiency, latency period and sporulation capacity — for 23 isolates representative of the most frequent genotype within each pathotype (106 314 0-G2 and 166 317 0-G1, identified by their combination of microsatellite markers). We tested these isolates on seedlings of Michigan Amber, a ‘naive’ wheat cultivar that has never been grown in the landscape, Apache, a ’neutral‘ cultivar with no selection effect on the landscape-pathotype pattern, and several cultivars that were frequently grown. We found that 106 314 0-G2 was more aggressive than 166 317 0-G1, with a consistency for the three components of aggressiveness. Our results show that aggressiveness plays a significant role in driving evolution in pathogen populations by acting as a selective advantage, even offsetting the disadvantage of lacking virulence towards a major Lr gene. Higher aggressiveness represents a competitive advantage that is likely even more pronounced when exhibited at the landscape scale as the expression of its multiple components is amplified by the polycyclic nature of epidemics.

## Introduction

Pathogenicity of plant pathogens is generally broken down into a qualitative term, ‘virulence’, and a quantitative term, ‘aggressiveness’ (Lannou, 2012). Virulence is the capacity of the pathogen to infect its host (compatibility) as opposed to avirulence, which expresses a resistance (incompatibility). This follows the gene-for-gene model involving a specific interaction between an avirulence gene (Avr) on the pathogen side and a corresponding qualitative resistance gene (R) in the plant (Flor, 1971; Dangl & Jones, 2001). In field conditions, R genes exert a high selection pressure, which contributes to shape pathogen populations. This pressure is all the higher as the R genes are deployed at high frequency in the landscape (Rouxel et al., 2003; Goyeau et al. 2006; Fontyn et al., 2022; Mundt, 2014; Rimbaud et al., 2018), favoring the selection of isolates carrying the corresponding virulences, hence resulting in a rapid decline of the immunity of the cultivated varieties. The aggressiveness describes both the parasitic fitness (Shaner et al., 1992) and the amount of damage caused to the host plant. It is formally defined as the quantitative variation of pathogenicity of a pathogen on its host plant (Pariaud et al., 2009a; Lannou, 2012) and is related to several life-history traits of the pathogen. Quantitative resistance, based in most cases on several quantitative trait loci or QTL (Niks et al., 2015), characterizes an incomplete immunity expressed by a reduction in symptoms intensity. The selection pressure exerted by quantitative resistance on aggressiveness components of pathogen populations is considered lower than that exerted by R genes (Mundt, 2014; Cowger & Brown, 2019).

The most widely assessed aggressiveness components for rust pathogens are infection efficiency, latency period and sporulation capacity (Pariaud et al., 2009a; Lannou, 2012; Azzimonti et al., 2013). Infection efficiency (IE) is the proportion of spores able to cause a new infection when deposited on the host plant tissues (Sache, 1997). Latency period (LP) is the length of time between the deposition of a spore and the appearance of most (usually 50%) of the sporulating structures (Parlevliet, 1975; Johnson, 1980; Pfender, 2001). Sporulation capacity (SP) is the number of spores produced per individual sporulation structure and per unit of time (Sache, 1997; Pariaud et al., 2009a).

Pathogen aggressiveness acts on the pathogen population dynamics, as it determines the rate at which a given intensity — incidence and severity — can be reached by a polycyclic disease (Azzimonti et al., 2022). Field experiments have shown that the most aggressive individuals tended to be selected over the course of an epidemic (e.g. Laloi et al., 2016; Suffert et al., 2018), highlighting that aggressiveness can be a significant component of the short-term evolution of pathogen populations. Several studies conducted under controlled conditions have suggested that the aggressiveness of fungal wheat pathogens — e.g. Puccinia graminis f. sp. avenae (Leonard, 1969) and Fusarium graminearum (Sakr, 2022) — increases after repeated cycles on the same host. Under controlled conditions, Lehman and Shaner (1997) have also demonstrated that the composition of a P. triticina population can be changed by selecting isolates with a shorter latent period after several cycles of asexual reproduction on a partially resistant cultivar. Additionally, Azzimonti et al. (2013) found that P. triticina isolates can adapt to quantitative resistance. The hypothesis that aggressiveness plays a role in the evolution of fungal plant pathogen populations is therefore based on experiments performed under controlled conditions with strains collected at large spatiotemporal scales and consistent with theoretical modelling approaches (e.g. Van den Berg et al., 2014; Rimbaud et al., 2018). So far, it has been rarely demonstrated experimentally at the pluriannual scale for real varietal landscape (e.g. Milus et al., 2009).

Leaf rust caused by Puccinia triticina is one of the most damaging wheat diseases, causing high yield losses worldwide (Huerta-Espino et al., 2011; Savary et al., 2019). Both qualitative and quantitative resistances play a role in the adaptive dynamics of P. triticina populations, which is broadly determined by the evolution of clonal lineages (Goyeau et al., 2007; Pariaud et al., 2009a; Kolmer, 2019; Zhang et al., 2020). Each P. triticina isolate can be characterized by its virulence profile, also known as a pathotype or race. Its combination of virulences can be identified at the seedling stage using a set of wheat lines carrying different Lr resistance genes. The deployment of a new qualitative resistance gene (Lr) — among the 82 that have been permanently designated in wheat so far (Bariana et al., 2022) — in the cultivar landscape is most of the time followed by the rapid emergence of pathotypes carrying the corresponding virulence, such as Lr28 in France, which has been overcome within two years after its introduction in the varietal landscape (Fontyn et al., 2022). This adaptive dynamic results in ‘boom-and-bust’ cycles of resistance deployment (McDonald & Linde, 2002).

Recently, Fontyn et al. (2022) showed that the domination of two P. triticina pathotypes in the French landscape during the decade 2006-2016 could not be fully explained by the deployment of Lr genes. Considering the Lr genes present in the varietal landscape from 2011 to 2015, the four most frequent Lr13, Lr37, Lr14a and Lr10 were overcome by both pathotypes 106 314 0 and 166 317 0. However, Lr3, the fifth most frequent Lr gene, should have limited the prevalence of the pathotype 106 314 0, avirulent to Lr3, as compared to 166 317 0, virulent to Lr3. Yet 106 314 0 remained at a frequency superior or equal to that of 166 317 0 until 2015. It was proposed that the longer than expected persistence of 106 314 0 in the landscape could be due to its higher aggressiveness. However, conclusively proving this assertion poses a formidable challenge, even for uredinologists accustomed to conducting experimental studies in quantitative epidemiology and having access to large collection of isolates.

Fontyn et al. (2023) reported that two isolates of the same pathotype are not necessarily of identical genotype, as defined by microsatellite markers, and therefore do not necessarily belong to the same clonal lineage, while identical genotypes may differ in one or several virulences. To underline this point, the authors proposed to use the term ‘pathogenotype’ to identify isolates based on a unique pathotype × genotype association. This analysis paved the way for two complementary approaches to investigate the hypothesis regarding the role of aggressiveness in the evolution of P. triticina populations. The first approach consisted in comparing, within each of the aforementioned dominant pathotypes 106 314 0 and 166 317 0, the ‘new’ pathogenotypes (106 314 0-G2 and 166 317 0-G2) with the ‘old’ ones (106 314 0-G1 and 166 317 0-G1, respectively), considering a large time period (2006-2016). With this strategy, Fontyn et al. (2023) found that the new pathogenotypes exhibited higher levels of aggressiveness compared to the old ones. They concluded that this fitness advantage can explain the replacement of isolates by others with the same virulence combinations. The second approach emerges as highly complementary, as it examines population evolution over a shorter timeframe: it involves comparing the aggressiveness of two pathogenotypes (106 314 0-G2 and 166 317 0-G1) that dominated from 2012 to 2015, and, unlike in the first approach, that differ not only in genotype but also in their virulence profiles. 106 314 0-G2 maintained high frequencies despite a significant disadvantage, namely being avirulent towards Lr3, a gene that was widely deployed at the time (Fontyn et al., 2022). The inception of this second approach propelled the current study, wherein the aggressiveness of 23 isolates representative of 106 314 0-G2 and 166 317 0-G1, was compared. Three aggressiveness components — IE, LP, and SP — were assessed on two cultivars considered ’neutral’ and ’naïve’, meaning that they have no selection effect on the cultivated landscape-pathotype pattern, and on five of the most commonly grown cultivars from 2006 to 2016.

## Material and methods

### Selection and purification of isolates

The 23 isolates used in this study were collected in 2012-2013 during annual surveys of P. triticina populations carried out throughout the wheat-growing areas in France over the last two decades (Goyeau et al., 2006; Fontyn et al., 2022). Eight isolates were selected from pathogenotype 106 314 0-G2 and 15 from pathogenotype 166 317 0-G1 (Table 1), which were identified as the most prevalent within each pathotype from 2012 to 2015 (Figures 4 and 5 in Fontyn et al., 2023), justifying their choice as case studies.

**Table 1.** List of the 23 Puccinia triticina isolates of the two pathogenotypes (106 314 0-G2 and 166 317 0-G1) collected in 2012-2013 and characterized for their agressiveness.

| Isolate | Pathotype * | Pathogenotype | Series | Cultivar sampled (Cs) |
| --- | --- | --- | --- | --- |
| BT13M108 | 106 314 0 | 106 314 0-G2 | 1 | Aubusson |
| BT13V026 | 106 314 0 | 106 314 0-G2 | 1 | Aubusson |
| BT13V143 | 106 314 0 | 106 314 0-G2 | 1 | Arezzo |
| BT12M024 | 106 314 0 | 106 314 0-G2 | 1 | Arezzo |
| BT12M302 | 106 314 0 | 106 314 0-G2 | 1 | Premio |
| BT12M293 | 166 317 0 | 166 317 0-G1 | 1 | Aubusson |
| BT12V078 | 166 317 0 | 166 317 0-G1 | 1 | Aubusson |
| BT13V066 | 166 317 0 | 166 317 0-G1 | 1 | Aubusson |
| BT13M239 | 166 317 0 | 166 317 0-G1 | 1 | Arezzo |
| BT13M259 | 166 317 0 | 166 317 0-G1 | 1 | Arezzo |
| BT12M042 | 166 317 0 | 166 317 0-G1 | 1 | Arezzo |
| BT12M371 | 166 317 0 | 166 317 0-G1 | 1 | Premio |
| BT12M264 | 166 317 0 | 166 317 0-G1 | 1 | Premio |
| BT12M079 | 166 317 0 | 166 317 0-G1 | 1 | Premio |
| BT12M033 | 106 314 0 | 106 314 0-G2 | 2 | Apache |
| BT12M391 | 166 317 0 | 166 317 0-G1 | 2 | Apache |
| BT12M192 | 166 317 0 | 166 317 0-G1 | 2 | Apache |
| BT12M326 | 106 314 0 | 106 314 0-G2 | 2+3 | Apache |
| BT12M321 | 166 317 0 | 166 317 0-G1 | 2+3 | Apache |
| BT13V189 | 106 314 0 | 106 314 0-G2 | 1+2+3 | Apache |
| BT12M202 | 166 317 0 | 166 317 0-G1 | 1+2+3 | Apache |
| BT12M378 | 166 317 0 | 166 317 0-G1 | 1+2+3 | Apache |
| BT12M109 | 166 317 0 | 166 317 0-G1 | 1+2+3 | Apache |

\* Seven-digit code adapted from the scoring system proposed by Gilmour (1973).

The recovery of these isolates was achieved as follows. Urediniospores were bulk-harvested from numerous infected leaves collected in a single field plot, purified from a single pustule and pathotyped on wheat differential lines at the seedling stage according to the method described in Goyeau et al. (2006). Among the bulks of urediniospores initially sampled in 2012 and 2013, 58 corresponded to the pathotype 106 314 0 (carrying virulences 1, 10, 13, 14a, 15, 17, 37) and 56 to the pathotype 166 317 0 (carrying virulences 1, 3, 3bg, 10, 13, 14a, 15, 17, 26, 17b, 37). One isolate from each of these 114 bulks stored at -80°C has been purified once again in 2020 and genotyped with 19 microsatellite markers (Duan et al., 2003; Szabo & Kolmer, 2007). Different genotypes were identified within each pathotype from these markers leading to the distinction of several pathogenotypes, i.e. different pathotype × genotype combinations (Fontyn et al., 2023). Indeed, identical genotypes may differ in one or several virulences, and, conversely, that different genotypes can have the same virulence profile while potentially expressing differences in their aggressiveness.

### Experimental design

Three aggressiveness components, infection efficiency (IE), latency period (LP) and sporulation capacity (SP), were assessed for the 23 selected isolates (Table 1). The 23 isolates were characterized in three successive series, according to the same protocol and under the same experimental conditions (Table 2). In series 1, we compared the aggressiveness of pathogenotypes 106 314 0-G2 and 166 317 0-G1 (collected on cultivars Apache, Arezzo, Aubusson and Premio) on two ‘neutral’ wheat varieties: (i) Apache, a commercial French cultivar shown in a previous study (Fontyn et al., 2022) to have no selection effect on the cultivated landscape-pathotype pattern, and (ii) Michigan Amber, considered intrinsically ‘naive’ both because it carries no known leaf rust resistance gene and because it has never been cultivated in France. In series 2 and 3, we compared the aggressiveness components of both pathogenotypes on five of the most frequently grown cultivars in the French landscape from 2006 to 2016: Aubusson, Premio, Sankara, Expert and Bermude. The isolates used in series 2 and 3 were collected on cultivars Apache and were tested on Aubusson and Premio (for series 2) and Bermude, Expert and Sankara (for series 3). To test the isolate × cultivar interaction in all series, eight wheat seedlings were used and the experiment was replicated three times, so that averages and median values were based on 24 data points.

**Table 2.** Experimental design for comparison of the aggressiveness of pathogenotypes 106 314 0-G2 and 166 317 0-G1.

| Series | Pathogenotype | Number of isolates | Sampled cultivar (Cs) | Tested cultivar (Ct) |
| --- | --- | --- | --- | --- |
| 1 | 106 314 0-G2 | 1 | Apache | Apache, Michigan Amber |
|  |  | 2 | Arezzo |  |
|  |  | 2 | Aubusson |  |
|  |  | 1 | Premio |  |
|  | 166 317 0-G1 | 3 | Apache |  |
|  |  | 3 | Arezzo |  |
|  |  | 3 | Aubusson |  |
|  |  | 3 | Premio |  |
| 2 | 106 314 0-G2 | 3 | Apache | Aubusson, Premio |
|  | 166 317 0-G1 | 6 |  |  |
| 3 | 106 314 0-G2 | 2 | Apache | Bermude, Expert, Sankara |
|  | 166 317 0-G1 | 4 |  |  |

### Evaluation of aggressiveness components

The pathotyping tests were performed in a greenhouse on eight-day-old wheat seedlings grown in a plastic box containing potting soil placed under standardized conditions as described in Fontyn et al. (2023). The inoculation was performed with 10 fresh (two-week-old) spores, picked one by one with a human eyelash under a dissecting microscope, then deposited on a 3 cm-long segment of the second leaf of each seedling, maintained with double-sided tape on a rigid plate coated with aluminum foil. The plants were then placed in a dew chamber. Uredinia were counted twice a day at 10-to 14-hour intervals, after the first uredinia break through the leaf epidermis and until no new uredinia appeared. Once the final number of uredinia had been reached, i.e. 9 days after inoculation, the spores that had already been produced were removed from the leaf with a small brush. Slightly incurved aluminum gutters (2 × 7cm) made from blind slats were positioned under each inoculated leaf (Figure 1) and all the newly produced spores after four days were removed by suction with a cyclone collector into a portion of plastic straw. Each portion of straw was weighed before and after spore harvesting. Infection efficiency (IE) was estimated for each leaf as the ratio between the final number of uredinia and the number of spores deposited (10 spores). Latency period (LP), expressed in degree-days based on the air temperature measured in the greenhouse every ten minutes, was estimated as the time between inoculation and the appearance of 50% of the total number of uredinia. Sporulation capacity (SP) was calculated by dividing the total weight of the spores collected from a single leaf by the number of uredinia on that leaf.

**Figure 1.**
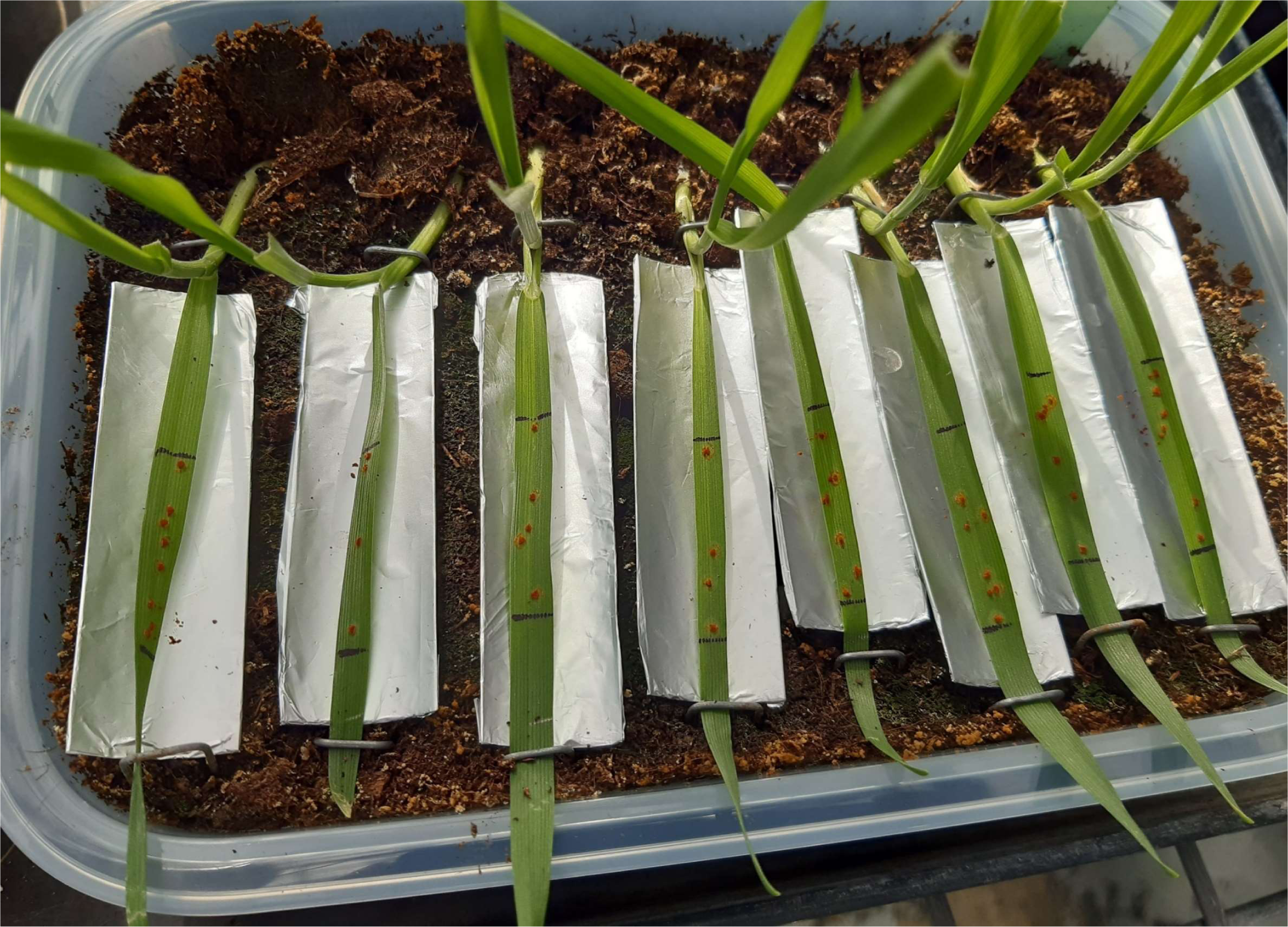
Experimental device consisting of incurved aluminium gutters positioned under wheat leaves presenting uredina, used to measure the sporulation capacity of each Puccinia triticina isolate.

### Statistical analyses

An ANOVA model (1) was used to examine each of the three aggressiveness components (the response variable Y), evaluated separately for each cultivar (Apache and Michigan Amber) in series 1. Our objective here was not to assess the effect of the cultivar since Michigan Amber was never grown in fields contrary to Apache. Thus, the ANOVA model was used to investigate each component of aggressiveness for each cultivar independently.

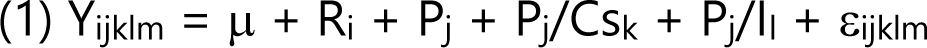

where Y_ijklm_ are the value of the aggressiveness component in replicate (R) i, of pathogenotype (P) j, sampled cultivar (Cs) k nested within pathogenotype, and isolate (I) I nested within pathogenotype. μ is the overall mean value for this trait and ε is the residual, representing the measurement error, with ε ∼ N(o, σ2).

A linear mixed model (2) was used to analyze the data of series 2 and 3 separately using the R package nlme (Pinheiro et al., 2017). The mixed model enables simultaneous multiple comparisons of two fixed effects, allowing for the analysis of the effect of each sampled cultivar on the two pathogenotypes.

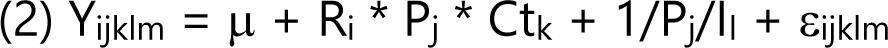

where Y_ijklm_ are the value of the aggressiveness component in replicate (R) i, of pathogenotype (P) j, tested cultivar (Ct) k, and isolate (I) I nested within pathogenotype.

R, P, Cs and Ct were fixed effects as well as the interaction between them, while I was a random effect. Significance for the fixed effect was calculated using the Satterthwaite method to estimate degrees of freedom and generate p-values for mixed models. Significance for the random effect was calculated based on the likelihood ratio chi-squared test. Post hoc test was performed using the R package emmeans (Lenth et al., 2018) for single effect, and the R package multcomp (Hothorn et al., 2016) for multiple comparisons of two fixed effects.

Log, √ or 1/x transformation was applied to IE, LP and SP when necessary, to obtain a normalized distribution of residuals. When the distribution of residuals could not be normalized by any transformation, a non-parametric Kruskal-Wallis test was performed to analyze the effect of genotype on the aggressiveness components. All the analyses were performed with R software version 4.1.0.

## Results

### Differences in aggressiveness between the two most frequent pathogenotypes, on neutral or naïve cultivars

When tested in series 1, pathogenotype 106 314 0-G2 appeared overall more aggressive than 166 317 0-G1. On the ‘neutral’ cultivar Apache, this difference was significant for latency period (LP) and sporulation capacity (SP) (Figure 2B and 2C; Table 3; p-values provided in Table 4). On the ‘naïve’ cultivar Michigan Amber, this difference was significant for infection efficiency (IE) and SP (Figure 2A and 2C). IE was higher for pathogenotype 106 314 0-G2 (69%) than for 166 317 0-G1 (64%) on Michigan Amber only. SP was higher for pathogenotype 106 314 0-G2 (0.092 mg.uredinium^-1^ on both Apache and Michigan Amber) than for 166 317 0-G1 (0.084 mg.uredinium^-1^ on Apache and 0.085 mg.uredinium^-1^ on Michigan Amber). LP was shorter for pathogenotype 106 314 0-G2 (135.5 degree-days) than for 166 317 0-G1 (141.5 degree-days) on cultivar Apache only.

**Figure 2.**
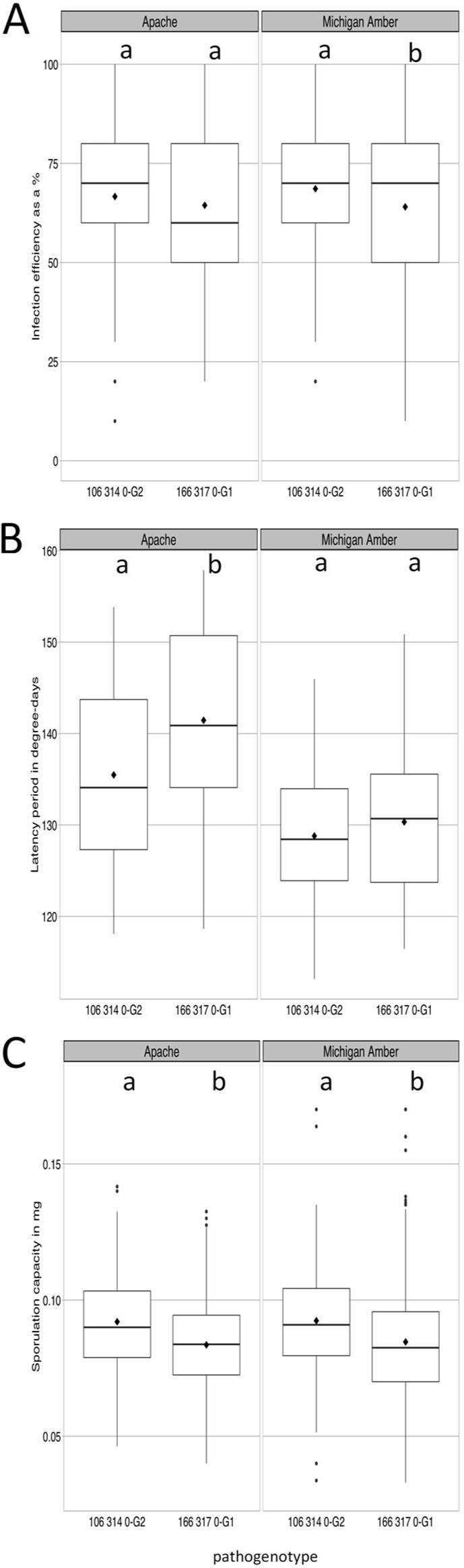
Comparison of three aggressiveness components of Puccinia triticina pathogenotypes 106 314-G2 and 166 317-G1 — infection efficiency in % (A), latency period in degree-days (B) and sporulation capacity in mg of spores.uredinium^-1^ (C) — measured on cultivars Apache and Michigan Amber. Within a box plot, black diamonds represent the mean value and horizontal bars the median value. Letters indicate statistical difference between pathogenotypes, resulting from Tukey test.

**Table 3.** Comparison of the aggressiveness components of the pathogenotypes 106 314 0-G2 and 166 317 0-G1 assessed on cultivars Apache and Michigan collected in 2012-2013 (series 1). Infection efficiency (IE) was measured as a %, latency period (LP) in degree-days, and sporulation capacity (SP) in mg of spores.uredinium^-1^.

| Pathogenotype | Isolate | Apache |  |  | Michigan Amber |  |  |
| --- | --- | --- | --- | --- | --- | --- | --- |
|  |  | LP | IE | SP | LP | IE | SP |
| 106 314 0-G2 | BT12M024 | 141.3 a | 69.6 ab | 0.087 a | 133.5 a | 76.4 a | 0.091 a |
|  | BT12M302 | 135.8 ab | 55.7 ab | 0.096 a | 127.2 bc | 63.7 ab | 0.082 a |
|  | BT13M108 | 133.7 ab | 72.5 ab | 0.090 a | 128.5 bc | 69.2 ab | 0.092 a |
|  | BT13V026 | 137.1 ab | 74.8 a | 0.092 a | 130.4 ab | 76.7 a | 0.100 a |
|  | BT13V143 | 133.7 ab | 57.4 b | 0.095 a | 128.5 bc | 60.0 b | 0.099 a |
|  | BT13V189 | 131.4 b | 70.4 ab | 0.091 a | 124.8 c | 64.8 ab | 0.092 a |
| 166 317 0-G1 | BT12M042 | 138.8 ab | 48.2 c | 0.090 a | 127.8 a | 56.5 cdef | 0.096 a |
|  | BT12M079 | 139.3 ab | 74.8 a | 0.079 ab | 131.2 a | 80.9 a | 0.086 a |
|  | BT12M109 | 140.9 ab | 67.3 ab | 0.087 a | 132.6 a | 70.0 abcd | 0.087 a |
|  | BT12M202 | 146.3 a | 72.7 a | 0.083 a | 132.1 a | 67.9 abcde | 0.089 a |
|  | BT12M264 | 141.7 ab | 63.8 abc | 0.081 ab | 130.1 a | 60.9 bcdef | 0.092 a |
|  | BT12M293 | 141.7 ab | 62.2 abc | 0.086 a | 132.4 a | 57.6 bcdef | 0.086 a |
|  | BT12M371 | 142.2 ab | 71.8 a | 0.080 ab | 128.1 a | 75.5 ab | 0.076 a |
|  | BT12M378 | 143.3 ab | 68.7 ab | 0.067 b | 129.7 a | 72.2 abc | 0.063 b |
|  | BT12V078 | 136.8 b | 66.7 ab | 0.091 a | 128.7 a | 50.0 ef | 0.087 a |
|  | BT13M239 | 143.4 ab | 59.1 abc | 0.082 a | 132.7 a | 53.68 def | 0.075 ab |
|  | BT13M259 | 142.6 ab | 52.9 bc | 0.088 a | 126.6 a | 48.18 f | 0.096 a |
|  | BT13V066 | 141.6 ab | 66.36 ab | 0.089 a | 132.6 a | 73.0 abc | 0.087 a |

Letters indicate a significant difference between isolates within the same pathogenotype based on a multiple-comparison test.

**Table 4.** P-values of the ANOVA model used to analyze the source of variations (R for replicate, P for pathogenotype, P/I for isolate and P/Cs for sampled cultivar) on the latency period (LP), infection efficiency (IE) and sporulation capacity (SP) in series 1.

| Tested cultivar | Aggressiveness component | R | P | P/I | P/Cs |
| --- | --- | --- | --- | --- | --- |
| Apache | LP | <0.0001 | <0.0001 | <0.0001 | 0.4 |
|  | IE | <0.0001 | 0.27 | <0.0001 | 0.6 |
|  | SP | <0.0001 | <0.0001 | 0.002 | 0.001 |
| Michigan | LP | <0.0001 | 0.01 | <0.0001 | 0.001 |
|  | IE | <0.0001 | 0.01 * | <0.0001 | <0.0001 |
|  | SP | <0.0001 | 0.0002 | <0.0001 | 0.001 |

\* significant interaction with replicate

Within each pathogenotype, we also evidenced a significant isolate effect on Apache and Michigan Amber (Table 4). Within pathogenotype 106 314 0-G2, the isolate effect was significant for LP and IE on both cultivars (Table 3). Within pathogenotype 166 317 0-G1, the isolate effect was significant for the three aggressiveness components on Apache, and for IE and SP only on Michigan Amber.

### Differences in aggressiveness between the two most frequent pathogenotypes, on five of the mostly grown cultivars

When tested on five of the mostly grown French cultivars in series 2 (Aubusson and Premio) and series 3 (Bermude, Expert and Sankara), pathogenotype 106 314 0-G2 appeared more aggressive than 166 317 0-G1 (Table 5), but the difference was statistically significant only for LP on Aubusson (135.5 vs 138.8 degree-days) and Sankara (140.7 vs. 143.9 degree-days). In addition, significant differences in varietal susceptibility were identified, with the effects of aggressiveness components showing consistent directionality. This consistency enhances the robustness of the conclusions drawn. In series 2, both pathogenotypes appeared more aggressive on Premio than on Aubusson (Table 5): 106 314 0-G2 had a significantly shorter LP and higher SP on Premio, and 166 317 0-G1 was significantly more aggressive on Premio than on Aubusson for all three components LP, IE and SP. In series 3, both pathogenotypes appeared more aggressive on Expert with higher IE and shorter LP as compared to Bermude and Sankara (Table 5). For both pathogenotypes, SP was not significantly different between Expert and Sankara, but was significantly lower on Bermude. Bermude was slightly less susceptible than Expert and Sankara to both pathogenotypes.

**Table 5.** Comparison of the aggressiveness components of the pathogenotypes 106 314 0-G2 and 166 317 0-G1 assessed on five of the most widely grown French cultivars during the 2006-2016 period. Infection efficiency (IE) was measured as a percentage, latency period (LP) in degree-days, and sporulation capacity (SP) in mg of spores.uredinium^-1^.

| Pathogenotype | Aggressiveness component | Series 2 |  | Series 3 |  |  |
| --- | --- | --- | --- | --- | --- | --- |
|  |  | Premio | Aubusson | Bermude | Expert | Sankara |
| 106 314 0-G2 | LP | 132.22 a | 135.35 b* | 142.09 b | 138.17 a | 140.74 b* |
|  | IE | 51.83 | 51.12 | 46.96 | 48.72 | 46.89 |
|  | SP | 0.134 a | 0.120 b | 0.121 a | 0.132 ab | 0.134 b |
| 166 317 0-G1 | LP | 131.23 a | 138.80 b* | 142.89 b | 138.44 a | 143.88 b* |
|  | IE | 54.04 a | 47.23 b | 41.94 a | 52.21 b | 45.53 a |
|  | SP | 0.127 a | 0.113 b | 0.119 a | 0.129 b | 0.132 b |

\* indicates a significant difference between the two pathogenotypes on the tested cultivar after a Tukey multiple-comparison test or a Wilcoxon-Mann-Whitney test.
Letters indicate a significant difference between cultivars within the same pathogenotype and series based on a multiple-comparison test.

## Discussion

### Despite a theoretical difference in fitness related to virulence to **Lr3**, two **P. triticina** pathogenotypes dominated similarly the cultivated landscape from 2012 to 2015

The two P. triticina pathotypes that extensively dominated the French landscape in the 2012-2015 period differed in their virulence profile with the presence of four more virulences in 166 317 0 than in 106 314 0, preventing the latter pathotype to infect varieties carrying the genes Lr3, Lr3bg, Lr17b and Lr26 (Fontyn et al., 2022). While Lr3bg, Lr17b and Lr26 are not present in French wheat cultivars, the frequency of Lr3 has continuously increased in the landscape since 2006, reaching 14% in 2014 (Fontyn et al., 2022). However, 166 317 0 was not more frequent than 106 314 0 from 2012 to 2014, as might have been expected (Papaïx et al., 2011; Kolmer, 2019; Zhang et al., 2020), provided that sampling involved cultivars with and without Lr3 at a quite similar frequency. This was the case in this study (see Tables 1 and 2 in Fontyn et al., 2022).

### The pathogenotype avirulent on **Lr3** exhibited higher aggressiveness, which should confer a significant selective advantage

In this study, we highlighted a significant difference in aggressiveness between two P. triticina pathogenotypes that not only differed in their genotype but also in their virulence profile, and co-dominated in the varietal landscape during a short time period (2012-2015). This result complements the findings obtained by Fontyn et al. (2023), which showed a difference in aggressiveness between ’new’ pathogenotypes and ’old’ ones, all of which had the same virulence profile, over a longer period of time (2006-2016). These two complementary approaches (see Introduction) provide a strong body of evidence attesting to the role of aggressiveness in the domination of certain isolates.

Significant differences in latency period (LP), infection efficiency (IE) and sporulation capacity (SP) were found, not only on the ‘neutral’ cultivar Apache and on the ‘naive’ cultivar Michigan, but also on Aubusson, Sankara, Premio, Bermude and Expert, for certain components at least. Although the difference between both pathogenotypes were not established for all the aggressiveness components, the ones that were significant all indicated that 106 314 0-G2 is more aggressive than 166 317 0-G1. Our results confirmed that most, but not all, of the aggressiveness components were related to each other in P. triticina (Azzimonti et al., 2022). LP, IE and SP result from the expression of an interaction between the pathogen and the plant, which is potentially driven by different QTLs (Azzimonti et al., 2014), and contribute to different extents to epidemic development. LP, which presents the most pronounced difference between 106 314 0-G2 and 166 317 0-G1, is likely the aggressiveness component having the strongest impact on epidemic dynamics since it drives the number of disease cycles during one epidemic season (Pringle & Taylor, 2002; Milus et al., 2006; Lannou, 2012). On cultivar Apache, the difference in LP was approximately half a day (11 h) at 15°C. As rust spores are known to be mainly dispersed towards the middle of the day (Pady et al., 1965), this difference may almost give a one-day advance in their dispersal if the environmental conditions (humidity, wind) are favorable. Each of the three aggressiveness components is supposed to confer significant, specific advantages to certain P. triticina isolates. The expression of these advantages in a diversified population in epidemic conditions deserves to be evaluated through pairwise or multiple competition experiments. Such competition experiments have been undertaken for P. striiformis f. sp. tritici (Bahri et al., 2009), subsequent to the demonstration that aggressiveness provides a selective advantage leading to pathotype replacement in stripe rust populations (Milus et al., 2009). To date, such investigations have not been conducted for P. triticina.

### The higher expression of aggressiveness in pathogenotype 106 314 0-G2 did not depend on the cultivar

Pathogenotype 166 317 0-G1 was never found more aggressive than 106 314 0-G2 whatever the cultivar on which it was tested and whatever the aggressiveness component considered. However, the significant differences in the aggressiveness of both pathogenotypes between the five cultivars tested in series 2 and 3 suggest that these cultivars differed in their susceptibility to P. triticina, consistently with the findings of Azzimonti et al. (2013). Pariaud et al. (2012) also evidenced aggressiveness variability between isolates within a given pathotype, despite the clonal reproduction of P. triticina in Europe. Specificity of quantitative resistance with regard to pathogen isolates, and thus the cultivar dependence of the aggressiveness expression without considering the effect of Lr genes, remains a matter of debate in the case of wheat leaf rust. Interactions between P. triticina isolates and wheat lines were found for latent period (Broers, 1989; Lehman & Shaner, 1996) and for sporulation capacity (Milus & Line, 1980), but Denissen (1991) did not find any specificity for these components. Later, Singh et al. (2011) stated that there was no isolate-specificity for quantitative resistance to the three wheat rust diseases in a large collection of CIMMYT breeding material.

### Higher aggressiveness expressed under controlled conditions may result in a significant competitive advantage when expressed at the landscape scale

Although Apache was considered as a ‘neutral’ cultivar and has a relatively high quantitative resistance level, its high proportion in the French landscape (> 8% over the decade 2003-2013) might have contributed to the selection of the most aggressive pathotype 106 314 0. Aubusson, Sankara, Premio, Bermude and Expert were the most frequently grown cultivars before 2012, and together they accounted for 11.1% of the cultivated landscape in 2012 (20.3% including Apache), 6.6% in 2013 (14.6% including Apache) and 3.0% in 2014 (9.0% including Apache). Therefore, the earliest pathotype 106 314 0 appears to be significantly advantaged over 166 317 0 on a significant part of the varietal landscape, and it is likely that the differences highlighted under controlled conditions have had a real impact in field conditions as the difference in aggressiveness might be expressed more intensely on adult plants than on seedlings (Milus and Line 1980). Knott (1991) found a higher (+25%) infection efficiency and a shorter (−4%) latent period of P. triticina on the upper wheat leaves than on the lower leaves, likely due to greater susceptibility of the upper leaves or to physiological effects.

It has already been shown that the P. triticina pathotype 073 100 0, which dominated the landscape between the late 1990s and early 2000s, had a higher aggressiveness on the cultivar Soissons as compared to other virulent pathotypes present in minor frequencies (Pariaud et al., 2009b). Soissons was the most commonly grown cultivar from 1991 to 1999, with a frequency in the French landscape going from 40.5% in 1993 to 15.3% in 1999. The domination of 073 100 0 was interpreted as a result of the adaptation of this pathotype to Soissons. Similarly, the advantage provided by the higher aggressiveness of 106 314 0-G2 on the mostly grown cultivars may explain the high frequency of 106 314 0 in the landscape in 2013-2014, despite the disadvantage conferred by its avirulence to Lr3.

Higher aggressiveness, when considered trait by trait (LP, IE, SP), may appear weak compared to the advantage conferred by a Lr gene. Indeed, the pathotype 106 314 0-G2 cannot infect at all wheat varieties carrying Lr3, which occupied up to 14% of the wheat surfaces in 2014. However, the smaller advantage of a few hours of shortened infection cycle (LP) combined with a few percent more released spores (SP) from each uredinia can be expressed across all other varieties, that is, 86% of the surface area. The frequency equilibrium between 106 314 0-G2 and 166 317 0-G1 observed in 2012, 2013 and 2014 suggests that a ‘moderate advantage’ expressed in a large majority of situations and a ‘strong advantage’ expressed in a small number of situations are equivalent.

It is important to approach our conclusions with caution as they were established based on a single case study. However, our findings represent a significant advancement in plant disease epidemiology as they corroborate assumptions and parameterizations utilized in theoretical models that emphasize the role of aggressiveness (e.g. Van den Berg et al., 2014; Rimbaud et al., 2018, 2021). These models suggest that the most promising trait for quantitative resistance is the latency period (LP), as it directly affects the number of epidemic cycles that the pathogen can complete during a given season. Our experimental results support this assumption. The significant differences observed between major pathogenotypes for other aggressiveness components will help to parameterize predictive models, particularly for leaf rust.

## Acknowledgments

We thank Nathalie Retout for her technical help in the preparation of the aggressiveness experiments. We thank the French Wheat Breeders groups ‘Recherches Génétiques Céréales’, ‘CETAC’, and ‘ARVALIS-Institut du Végétal’ for their help in collecting strains of P. triticina from their nurseries and field trials.

## Funding

This research was supported by a PhD fellowship from the INRAE department ‘Santé des Plantes et Environnement’ (SPE) and from the French Ministry of Education and Research (MESRI) awarded to Cécilia Fontyn for the 2018-2022 period. It was also supported by several French FSOV (‘Fonds de Soutien à l’Obtention Végétale’) grants FSOV 2004 K, 2008 G, 2012 Q), and by the European Commission, Research and Innovation under the Horizon 2020 program (RUSTWATCH 2018-2022, Grant Agreement no. 773311-2). INRAE BIOGER benefits from the support of Saclay Plant Sciences-SPS (ANR-17-EUR-0007).

## Data Availability Statement

Data, scripts, and code are available in the INRAE Dataverse online data repository at https://doi.org/10.57745/QDF06W

## Conflicts of Interest

The authors declare that the research was conducted in the absence of any commercial or financial relationships that could be construed as a potential conflict of interest.

## Notes

https://doi.org/10.57745/QDF06W

### Competing Interest Statement

The authors have declared no competing interest.

### Summary of Updates

Plant resistances impose strong selective pressure on plant pathogen populations through the deployment of resistance genes, which leads to the emergence of new virulences. The pathogen adaptation also involves other parasitic fitness traits, especially aggressiveness components. A previous study on Puccinia triticina, the causal agent of wheat leaf rust, revealed that the distribution frequency of virulences in the French pathogen population cannot be fully explained by the major resistance genes deployed in the landscape. From 2012 to 2015, two dominant pathotypes (distinguished by their combination of virulences) were equally frequent despite the theoretical advantage conferred to one pathotype (166 317 0) by its virulence to Lr3, frequent in the cultivated landscape, whereas the other (106 314 0) is avirulent to this gene. To explain this apparent contradiction, we assessed three components of aggressiveness - infection efficiency, latency period and sporulation capacity - for 23 isolates representative of the most frequent genotype within each pathotype (106 314 0-G2 and 166 317 0-G1, identified by their combination of microsatellite markers). We tested these isolates on seedlings of Michigan Amber, a 'naive' wheat cultivar that has never been grown in the landscape, Apache, a 'neutral' cultivar with no selection effect on the landscape-pathotype pattern, and several cultivars that were frequently grown. We found that 106 314 0-G2 was more aggressive than 166 317 0-G1, with a consistency for the three components of aggressiveness. Our results show that aggressiveness plays a significant role in driving evolution in pathogen populations by acting as a selective advantage, even offsetting the disadvantage of lacking virulence towards a major Lr gene. Higher aggressiveness represents a competitive advantage that is likely even more pronounced when exhibited at the landscape scale as the expression of its multiple components is amplified by the polycyclic nature of epidemics.

